# Size dependent regeneration capacity of functionalized Capra ear derived micro-tissue scaffolds for treatment of cartilage defects

**DOI:** 10.1101/2021.08.10.455755

**Authors:** Pritiprasanna Maity, Sumanta Mukherjee, Subahayan Das, Mahitosh Mandal, Santanu Dhara

**Affiliations:** School of Medical Science and Technology, IIT Kharagpur, Kharagpur, India 721302; Production Engineering Department, BIT Sindri, Dhanbad, Jharkhand, India 828123

**Keywords:** Capra ear, Collagen type II, Micro-scaffold, Cartilage regeneration

## Abstract

Cartilage regeneration remains a great challenge in orthopedic treatment owing to their avascular in nature and lack of self-healing ability. Various clinical treatment options are widely used including osteochondral graft transplantation, micro-fracture, blood clot formation and tissue debridement to recover the damaged cartilage. In this circumstance, cartilage defect recovery via tissue engineered functional micro tissue delivery is becoming an emerging trend in musculoskeletal therapeutics. In this study, functional micro-scaffolds (MS) were generated from *Capra* ear cartilage, and were separated into size-wise groups. The scaffolds were decellularized via NaOH treatment. The cell adhesion study indicated that *Capra* adipose tissue derived mesenchymal stem cells (ADMSCs) adhesion is more in ~100 μm MSs in comparison to 150-300 μm MSs. It may be assumed that the cells are compatible to grow on fibrous surface area (100 μm) in comparison to dense surface (150-300 μm). Further, 100 μm MSs were transformed into functional micro tissues (FMTs) in presence of high density ADMSCs in a hanging droplet culture system. The FMTs was transferred to F127 block polymer hydrogel for 3D culture. After 21 d cultures, the FMT clusters were evaluated for quantitative gene expression. To assess the *in vivo* cartilage defect regeneration potential, FMTs were delivered to rabbit auricular cartilage defect for 15, 30 and 60 d studies. The H&E-stained histological analysis showed that the cartilage defect is almost healed in 60 d study in comparison to 15 and 30 d study.

## 1. Introduction

Interest in ex-vivo organoid models is rapidly growing for various therapeutic applications. Owing to their native tissue-like properties, *ex vivo* organoid model has immense utility in reconstructing critical defects as promising graft source [1]. The recent development of ex-vivo organoid model involves both matrix-based as well as matrix free culture. While considering matrix based culture, the decellularized tissue could be explored as vehicle for delivering drugs, growth factors and cells [2].

The existing techniques for assessment of in-vivo tissue regeneration are mainly based on either in-vitro models or animal studies. Therefore, it is very difficult to correlate findings between studies investigated within an in-vitro microenvironment and complex in vivo implantation in physiological system, where complex cell-material interaction is predominant [3]. Moreover, for detail understanding of molecular cross-talk between cell and material, a large number of animals must be sacrificed that limits widespread application of existing in vivo model-based screening techniques for data validation.

To address such challenges, ex-vivo organoid culture based theranostic approaches have been developed in recent years for studying organoids either as potential drug screening model or substituting tissue transplantation for recovery of defects. Engineered functional micro tissues have immense potential in providing native tissue micro-environment within defects, stimulating secretion of appropriate bio-chemical cues as well as accelerating cell migration towards faster wound healing [4].

Cartilage is one of the important connective tissues existing in body parts, such as ear, nose, rib cage, intervertebral discs, bronchial tubes, meniscus and bone joints. Articular cartilage effectively distributes excessive physiological stress to the subchondral bone and also provides an extremely low friction surface to the joints during articulation [5]. However, lesions in articular cartilage may provide abnormality in joint function by preventing their self-healing ability and finally leading towards osteoarthritis [6]. Although, cartilage regeneration is one of the major concerns in orthopedic treatment due to their avascularity and limited self-reparative ability can reasonably delay complete reconstruction of cartilage defects or prevention of further deterioration.

Considering such limitations, engineered micro-scaffolds has advantageous for regeneration of defected cartilage. Moreover, after transformation of micro-scaffolds to its functional stage by culturing with stem cells, particles can undergo cartilaginous differentiation in absence of exogenous factor [4]. The functional micro-scaffolds and their aggregates can be directly used for cartilage regeneration or delivered by a hydrogel for sufficient sticking to the wound bed [7].

Here, we developed a method for effective separation of various sized micro-scaffolds from pulverized tissue and cartilage organoid by hanging culture in fabricated device. Primarily micro-scaffolds were decellularized using NaOH solution followed by alcoholic separation based on different sizes, ranging from 100 μm to 300 μm. ADMSCs were cultured on those scaffolds for subsequent spheroid formation.

The spheroids were evaluated by live-dead, ex-vivo cartilaginous differentiation and gene expression analysis. To understand the significance of the role of small particles in effective organoid formation, a plausible model was drawn for identifying the cell-material interaction. Further regenerative potential of organoids was assessed through in vivo implantation in rabbit auricular cartilage (Fig. 1).

**Fig. 1.**
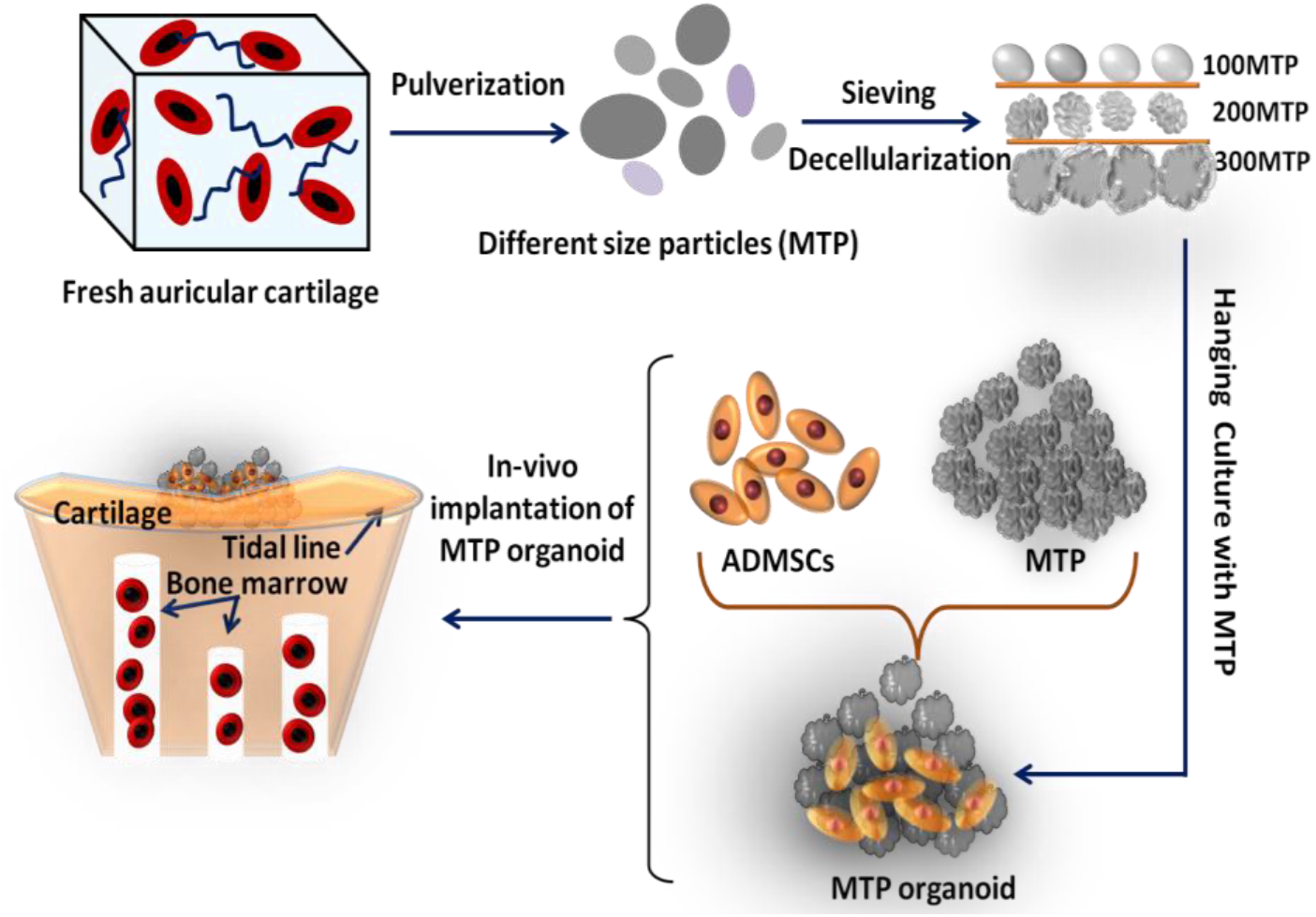
Schematic representation of micro-scaffold isolation, separation, spheroid formation and application to cartilage defects.

**Fig. 1.**
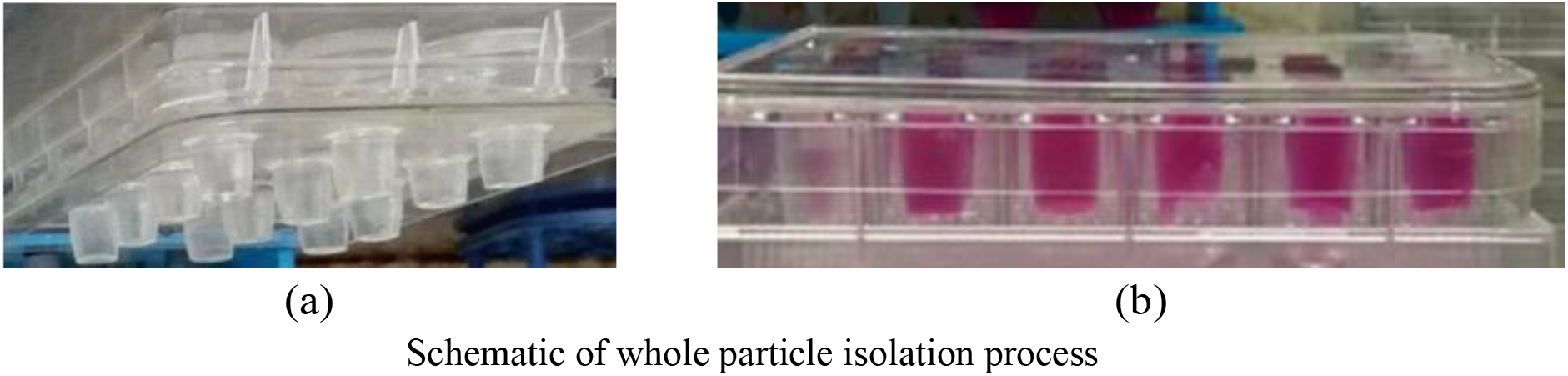
(a) Trimmed parts of the Eppendorf tubes attached to the lid of cell culture plate, and (b) Hanging drop culture in the setup

## 2. Materials and methods

### 2.1 Isolation, decellularization and characterization of micro scaffolds derived from *Capra* ear cartilage

*Capra* ears, freshly sourced from a local slaughter house, were thoroughly washed and mechanically pulverized after peeling off the skins. For decellularization, the tissues were treated with 0.1-0.3 M NaOH in PBS for 6 h and washed with PBS with the pH adjusted to neutral range. Decellularized tissues were further treated with 50 U/ml DNase I and 1 U/ml RNase (Sigma- Aldrich) for 4 h at 37 °C to remove the traces of DNA and RNA, respectively. Finally, the decellularized tissues were washed with PBS, lyopholized and immediately crushed to obtain the micro-scaffolds.

To separate the scaffolds into size-wise groups, the crushed tissue was vortexed with 70% alcohol for different time lengths. The particles precipitated within 60 s post 30 s and 60 s of vortexing were collected as the first the second group, respectively, and the particles precipitated within 60 s from the supernatant were collected as the third group. 60 s vortexing, 60 s precipitation: supernatant

For histological analysis, the decellularized particles were fixed with 4% paraformaldehyde followed by PBS wash and dehydration by alcohol gradation (70%, 90% and 100 % alcohol). The dehydrated particles were embedded in paraffin, stained with H&E, and sectioned with microtome for further analysis.

sGAG quantification was carried out by Alcian blue method, as reported earlier [8]. For that purpose, native and decellularized MSs (1 mg/ml) were digested with 0.1 M PBS (pH 6.8) containing 10 mM cysteine hydrochloride (Sigma, USA), 125 μg/ml papain (Sigma, USA) and 2 mM Na2EDTA (Sigma, USA) at 60 ºC for 24 h. The total sGAG content was measured using Alcian Blue 8GS (SRL, India). Standard curve was developed using different concentrations (12.5, 25, 50, 100, 200, 400, 600, 800 μg/ml) of chondroitin sulfate. The absorbance of the Alcian blue-sGAG complex was measured at 595 nm wavelength using iMarkt microplate reader and sGAG contents were estimated from the standard curve.

### 2.2 Morphology analysis

Morphologies of the MSs were analyzed from optical as well FESEM (EVO 60, Carl Zeiss, Germany) micrographs using ImageJ. Micro-CT (GE phoenix v|tome|x) was also performed with kV and current to analyze the morphologies from 3D reconstructions.

### 2.3 Preparation of Pluronic F 127 hydrogel

Pluronic F127 copolymer (20%) was dissolved in DMEM media and transferred to refrigerator for complete dissolution. The sol-gel mix was shaken vigorously and maintained at 4 °C overnight. The hydrogel, transformed to a clear translucent dispersion after several freeze-thaw cycles, was used for the study [9].

### 2.4 Hanging droplet culture for spheroid formation

Formation of spheroid-like constructs from the micro-scaffolds was studied by conducting hanging droplet culture of micro-scaffolds enriched with ADMSCs in a simple and frugal culture system developed in-house. For construction of the hanging culture system, the cap and 4 mm of the conical section were removed from 0.5 ml Eppendorf tubes, and modified tubes were attached to the inside of the lid of 12 well cell culture plate using PDMS (Fig. 2(a-b)). Micro-scaffolds were sterilized with 70% alcohol followed by washing with 1% antibiotic-gentamicine solution in PBS and incubated with complete DMEM media for overnight at 37 °C CO_2_ incubator. All the three types of scaffolds were separately seeded with ADMSCs isolated from *Capra* adipose tissue following a protocol reported earlier [9], and were placed in the hanging culture system. While placing the lid, some amount of PBS with antibiotic-antimicotic (1X) solution was introduced in all of the wells to maintain the humidity in the well. Post 6 days culture under gravitational force, the micro scaffold constructs were transferred to F127 hydrogel for 21 days culture. During the culture, the media was replaced every 3^rd^ day.

### 2.5 Live-Dead assay

Live-Dead assay was performed using Live–Dead staining kit (Invitrogen, USA). The ADMSCs-MS constructs were incubated for 30 min at 37 °C temperature in a solution containing 2 μM calcein aceto-methoxy (AM) and 4 μM ethidium homodimer. Fluorescence micrographs of the incubated constructs were captured using fluorescent microscope (Carl Zeiss, Germany) with excitation filters of 450–490 nm (green, Calcein AM) and 510–560 nm (red, ETD-1) using ZEN software.

### 2.6 Gene expression analysis

21 d post seeding, RNA was harvested from the micro tissue constructs using RNA extraction kit (Invitrogen, India). Synthesis of cDNA and PCR amplification were carried out (Thermo Scientific, USA) using primers enlisted in Table 2 to evaluate the expressions of chondrogenesis related genes, namely, collagen type II (COL II), aggrecan (CAN), and transcription factor SOX-9.

**Table 2.**
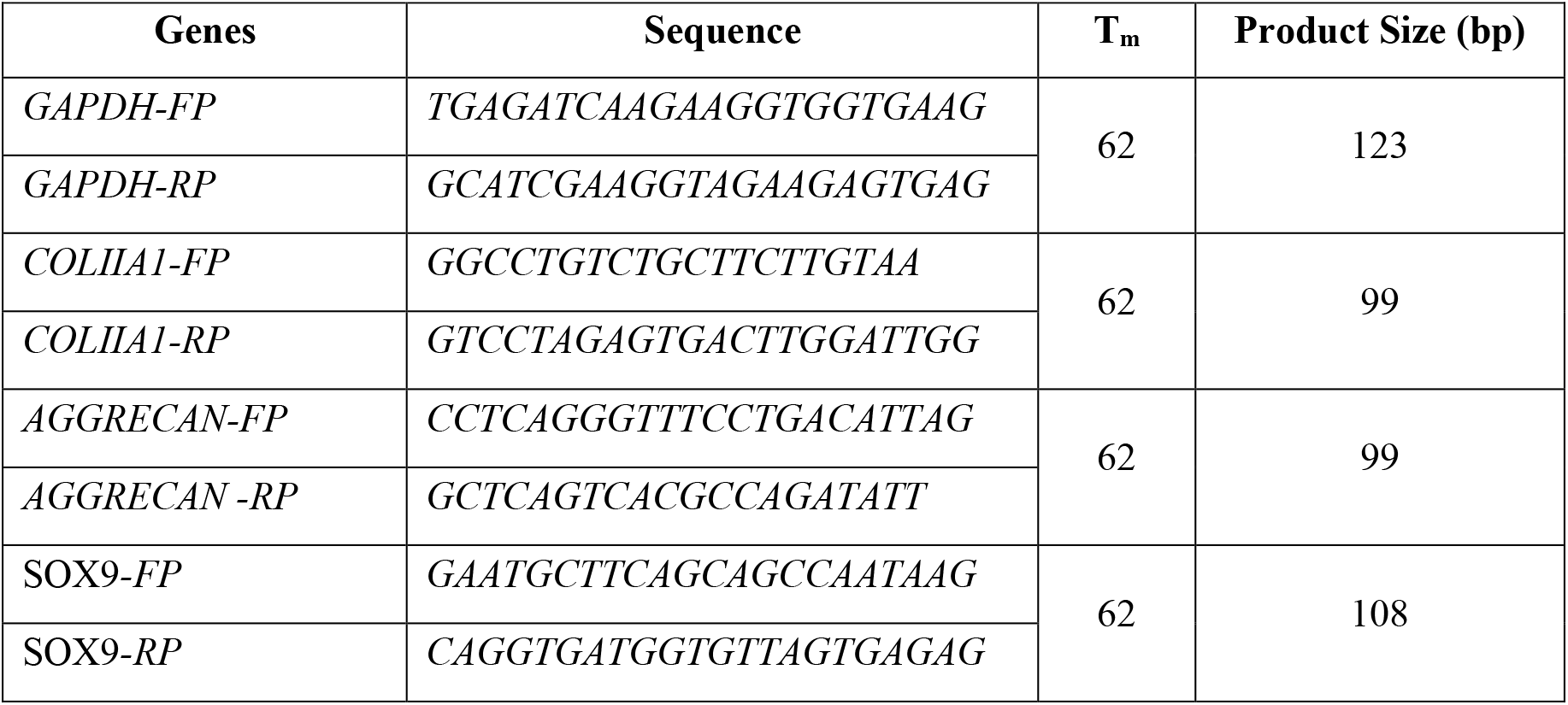
Primers used for qRT-PCR analysis of chondrogenic potential of micro tissue constructs.

### 2.7 In-vivo study in rabbit ear cartilage defect model

Efficacy of the micro-scaffolds towards regeneration of auricular cartilage defects was assessed in rabbit model using three New Zealand rabbits (weight 2.5–3.0 kg). All the rabbits were anesthetized with Midazolam (0.2 ml) followed by local anesthesia of the selected zone for surgery, and after skin hair removal, two 7-9 mm defects were generated in inner side of one auricle each rabbit, leaving the other auricle intact. MS-F127 hydrogel was delivered to one of the defects for each rabbit, keeping the other as control. Regeneration of the tissues was evaluated 15 d, 30 d and 60 d post-surgery. On the 60th day, the tissue samples were collected to further analysis after sacrificing the animals. Necessary approval for *in-vivo* studies in rabbit model was obtained from the Institutional Animal Ethical Committee of Indian Institute Technology, Kharagpur (IIT Kharagpur), India before initiating the study.

### 2.8 Assessment of cartilage regeneration

After sacrificing the rabbits, the ears were collected and washed with PBS and immediately transferred to micro-CT (GE phoenix v|tome|x) for tissue ingrowth analysis. The tissues were scanned with X-ray radiation (85 kV voltage, 75 μA beam current) to obtain 3D reconstructions of the defect zones. for. Quantitative evaluation of neo-cartilage tissue formation was performed using VG Studio MAX (Volume Graphics) software.

For histology, tissues were fixed with 4% paraformaldehyde, dehydrated with alcohol gradation (70%, 90%, 100%, and xylene fixation) followed by paraffin embedding and tissue sectioning. The sections were stained with Hematoxylin & Eosin (H&E), Masson Trichrome (MT) staining, and imaged with optical microscope (Car Zeiss, Germany).

### 2.9 Statistical analyses

All the experiments except the animal study were carried out in triplicate and the results are represented as means ± standard deviation. Origin software (Version: 8.5) was utilized for the statistical analyses. The significance of the experimental and control groups was calculated using analysis of variance (ANOVA) and P < 0.05 was considered statistically significant.

## 3. Results

### 3.1 Isolation and decellularization of *Capra* ear cartilage

Decellularization of the tissue was performed in 0.1-0.3 M NaOH/PBS solutions. H&E staining showed existence of cells in tissue treated with 0.1 M NaOH (Fig. 3(a)), whereas MSs treated with 0.2 M and 0.3 M NaOH were completely devoid of cells and nuclei (Fig. 3(b-c)). However, 0.3 M NaOH treatment resulted in distortion of native tissue structure, and accordingly, 0.2 M NaOH treatment for 6 h was selected as the optimum condition for decellularization of the tissue. Similarly, different concentrations of alcohol were used to separate the scaffolds by size, and 70% ethanol was found to be most effective for the separation by vortexing.

**Fig. 3.**
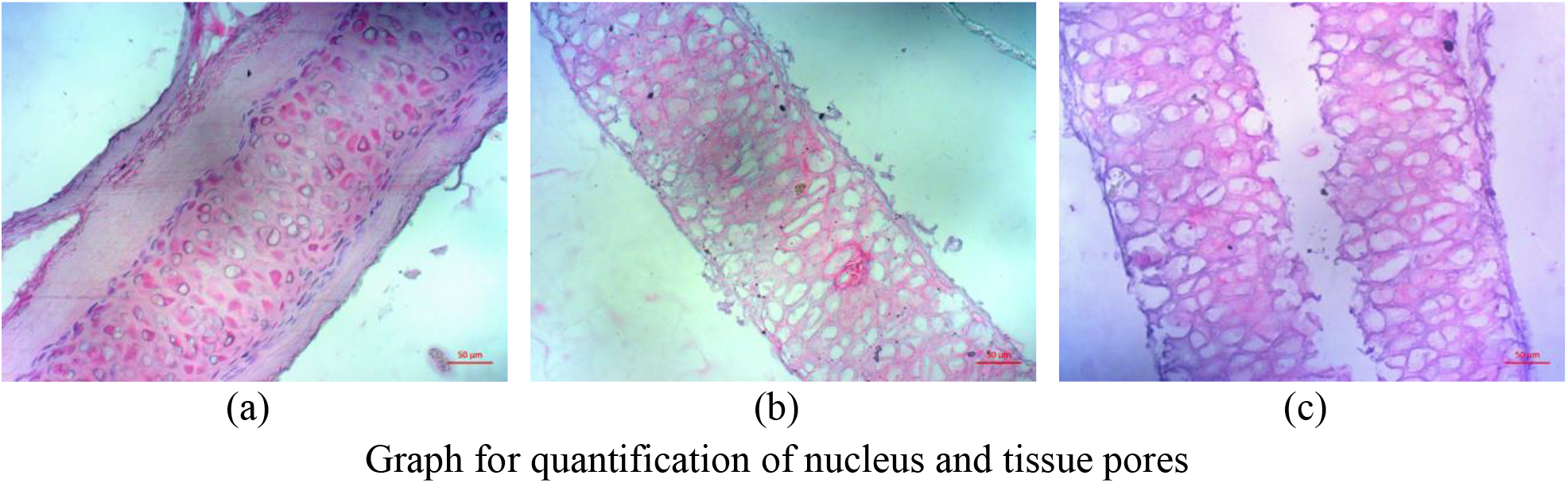
Decellularized tissue: (a-c) Treated with 0.1 M, 0.2 M, and 0.3 M NaOH, respectively.

sGAG assay showed that although the quantity of sGAG did not significantly change from the native tissue when the tissues were treated with 0.1 M NaOH for decellularization, increasing the concentration of NaOH deteriorated the quantity of sGAG (Fig. 4).

**Fig. 4.**
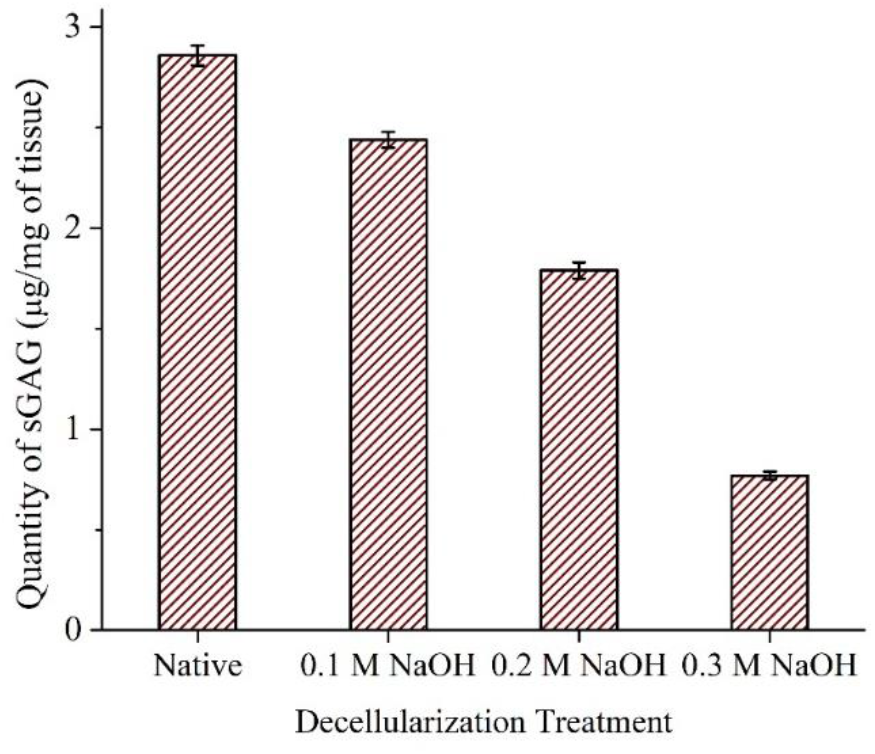
Quantification of sGAG present in different decellularized MSs.

### 3.2 Characterization of micro scaffolds

The FESEM micrographs evidenced that the process generated fibrous micro scaffolds with different sizes, and the vortexing-based separation process was able to segregate the micro scaffolds into size-wise groups (Fig. 5). The scaffolds separated after 30 s of vertexing were the largest in size, having an average size of 299.68 ±74.11 μm, while the scaffolds precipitated after 60 s of vortexing and the scaffolds precipitated from the supernatant were 152.0 ± 23.1 μm and 95.82±24.36 μm in size, respectively (Table 4). For ease of identification, these groups will be referred to as 300 μm MS, 150 μm MS, and 100 μm MS, respectively.

**Fig. 5.**
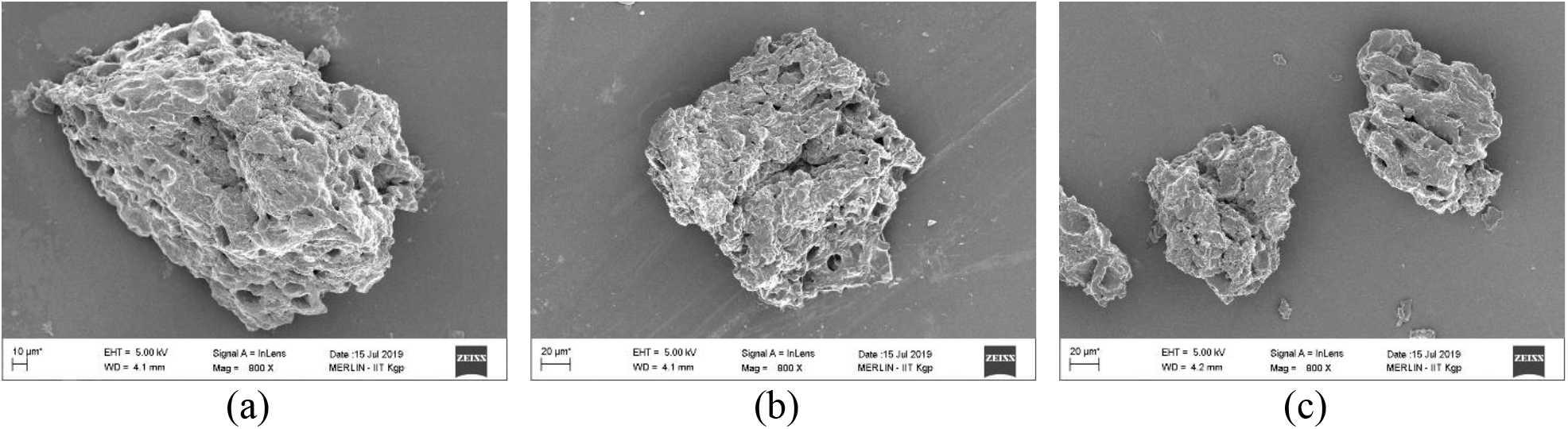
FESEM images of scaffolds: (a) 300 μM MS, (b) 150 μM MS, and (c) 200 μM MS.

**Table: 4.** Size dependent MS separation by ethanol at different time length.

| Micro scaffold type | Size (d) ± SD ($\mu\text{m}$) | | Circularity ± SD (0-1) |
| --- | --- | --- | --- |
| Gr. 1 (300 $\mu\text{m}$ MS) | $299.68 \pm 74.11$ | 225-375 | $0.51 \pm 0.29$ |
| Gr. 2 (150 $\mu\text{m}$ MS) | $152.0 \pm 23.1$ | 125-175 | $0.59 \pm 0.25$ |
| Gr. 3 (100 $\mu\text{m}$ MS) | $95.82 \pm 24.36$ | 75-125 | $0.33 \pm 0.22$ |

Close observation of the scaffolds revealed the microporous nature of the scaffolds resulting from the network of collagen fibrils (Fig. 6(a)), and the characteristic D-banding of collagen fibrils was also visible (Fig. 6(b)).

**Fig. 6.**
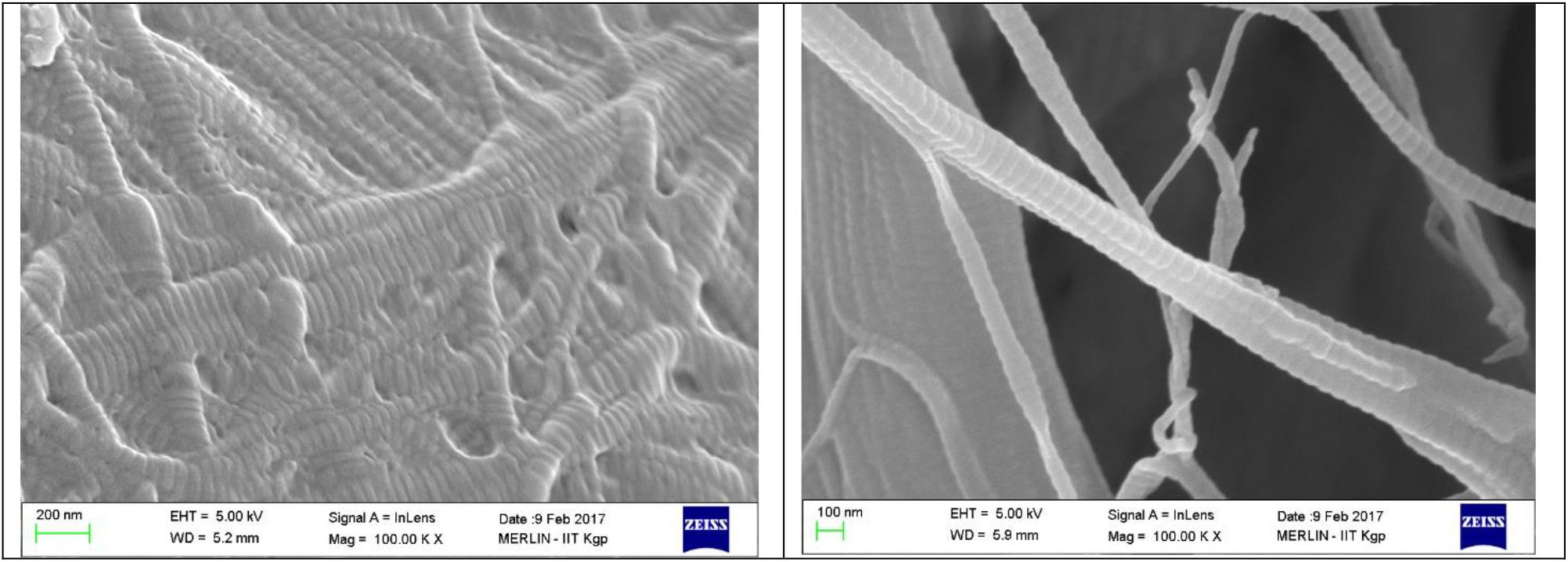
Architecture of the scaffolds: (a) Fibrous network of collagen fibrils, and (b) Characteristic D-banding.

**Fig. 7.**
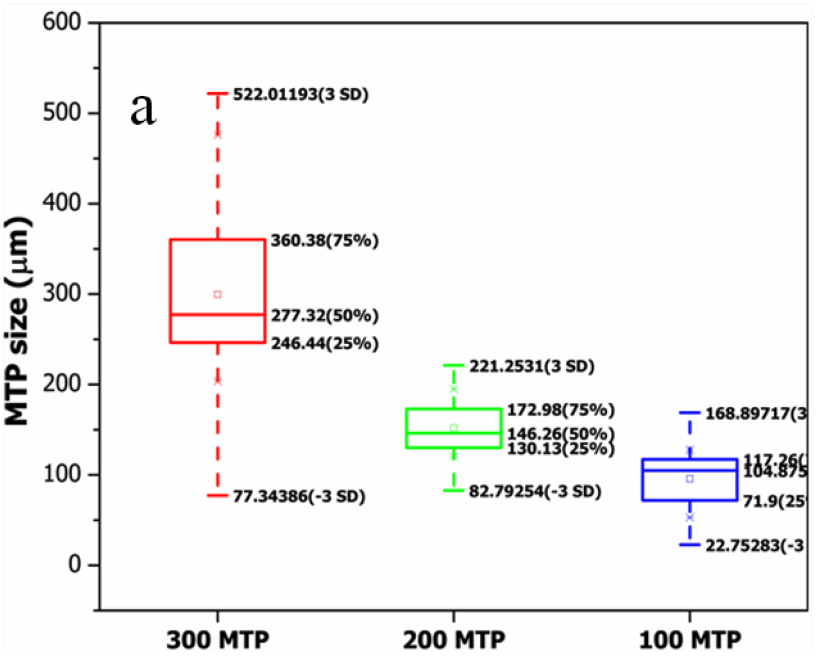
Quantitative analysis of different groups of micro-scaffolds (300, 150 & 100 μm).

### 3.3 Hanging culture for ex-vivo spheroid formation

For chondrogenic differentiation, ex-vivo spheroid culture method [4] was adopted using the fabricated device. The popular method for preparation of micro tissue organoid is drop casting on the plate with relatively less amount of media, which restricts duration of culture (Fig. 8(a)). The high density ADMSCs culture was performed in our fabricated device (Fig. 8(b-c)) for longer time using higher volume of media. MS constructs transformed into micro tissue after 3-4 days hanging culture. Cell adhesion and distinct micro tissue formation was observed in 100 μm MS (Fig. 8(f)) where 150 and 300 μm MS showed loose particles with less micro tissue formation (Fig. 8(d-e)).

**Fig. 8.**
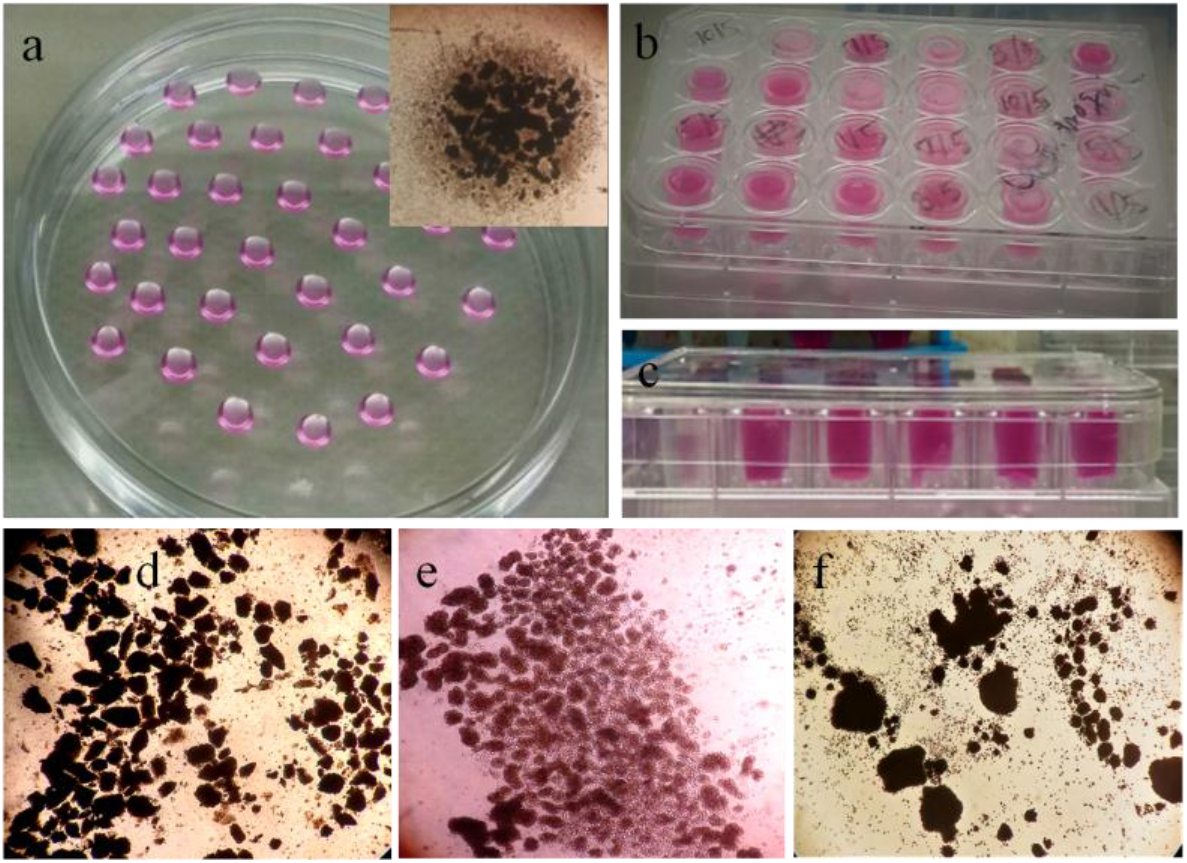
(a) Hanging culture performed on conventional drop cast method, (b-c) small piece of eppendrof attached on 6 well plate lids for prolonged hanging culture and (d-f) cell adhesion on 300, 150 and 100 MS.

### 3.4 Live/Dead imaging

The live/dead assay indicated that ADMSCs were attached on the surface of all the MSs after 3 d culture (Fig. 9 (a-c)). However, the cells were more prone to be attached on surface of 100 μm MSs (Fig.11 a) as compare to 150 μm and 300 μm MSs (Fig.11 b-c). Cellular aggregates were also observable on the surface of 100 μm MSs whereas larger particles showed limited cell adhesion. No dead cell was visible on any of the particles.

**Fig. 9.**
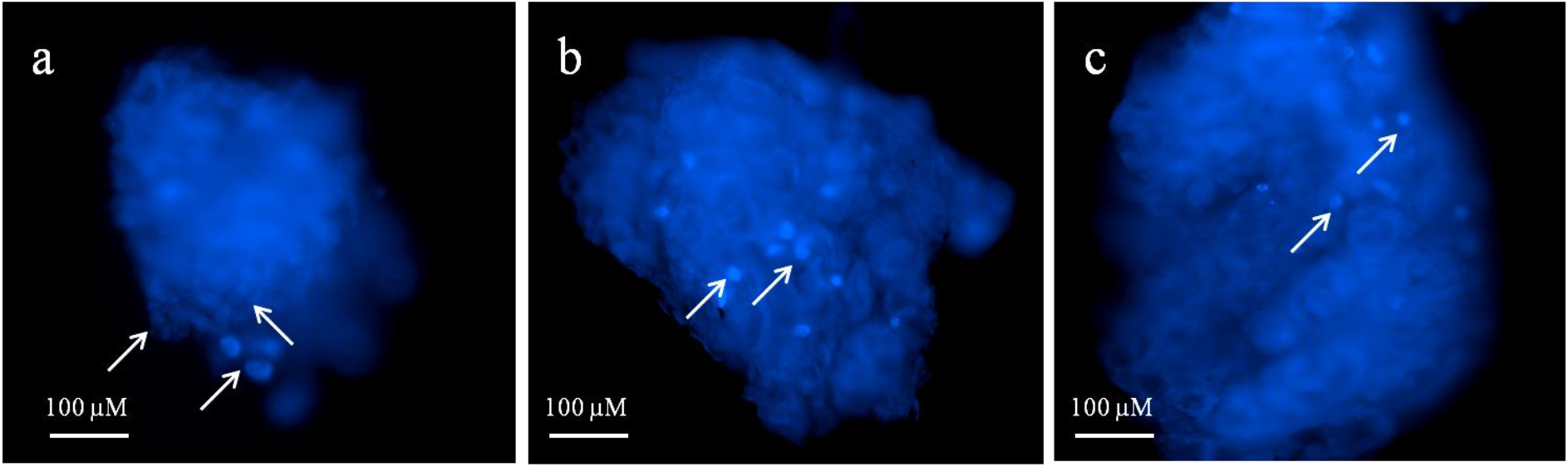
Live-dead assay of different size MS at 3 d hanging culture. (a) Formation of cell aggregates on 100 μm MS surface, (b) Less number of cell aggregates observed on 200 μm MS surface and (c) Fewer cells attached on the surface of 300 μm MS.

### 3.5 Chondrogenic differentiation study

Gene expressions were analyzed to quantify the mRNA expression profile of chondrogenic marker in the cellular clusters formed on the MSs. It was evident from the qRT-PCR data that expression of the chondrogenic markers (COL II, SOX 9 and Aggrecan) were significantly up-regulated for 100 μm MS organoid in comparison with 200 μm and 300 μm MS organoid after 21 d culture (Fig. 10). For the 100 μm MS organoids, the expressions of COL II, SOX 9 and Aggrecan were 4, 3, and 8 times, respectively, as compared to GAPDH, whereas, for the 150 μm MS organoid and the 150 μm MS organoids, the expressions of COL II, SOX 9 and Aggrecan were 0.64, 0.44, 0.77 times and 1.01, 1.04,1.35 times, respectively.

**Fig. 10.**
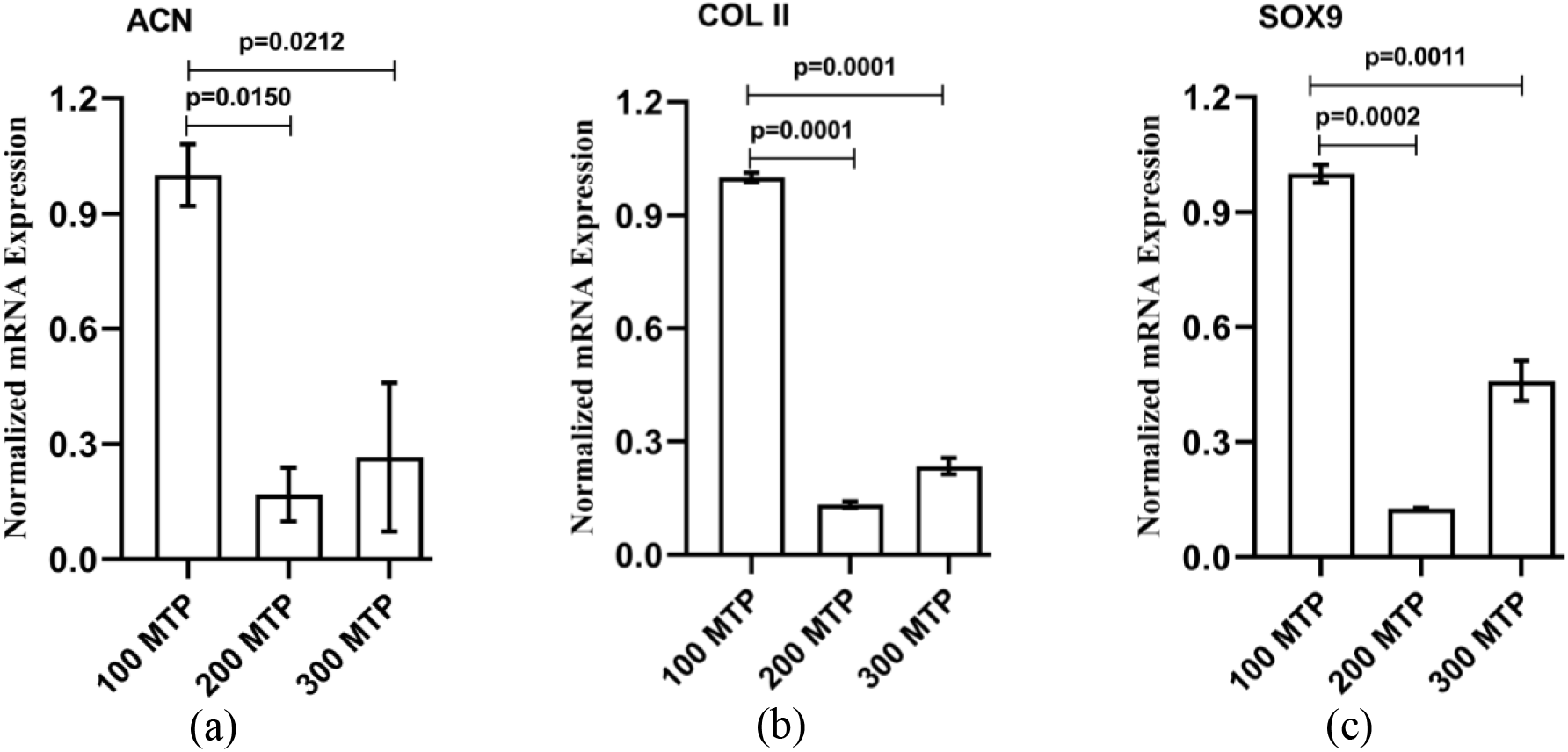
Relative quantification of gene expressions of different-sized micro-scaffolds

### 3.6 *In-vivo* rabbit auricular cartilage defect healing using 100 μm MS organoid

#### Gross observation of auricular cartilage defect

From gross observation, the F127 hydrogel incorporated 100 μm MS organoid was highly adjusted to native auricular cartilage tissue with no signs of infection or graft rejection at the defect area. The defect area was found to be gradually closing, initially with thick skin, and then by cartilage tissue growth. Initiation of cartilage tissue formation was observable 30 d post-surgery, and the defect area was completely recovered after 60 days.

#### Non-invasive tissue in growth assessment by Micro-CT imaging

Micro-CT scanning was carried out to assess the neo-tissue formation in cartilage defects. The analysis results showed that defect healing after 100 MTP organoid transplantation was gradually increased in accordance with time length. After 15 d and 30 d, auricular cartilage defect regeneration was 31.60±2.78 % and 40.56± 1.21 % higher in respect to void area. After 60 d, the defected cartilage was almost healed (81.92±5.63 %) compared to void area (Fig. 11).

**Fig. 11.**
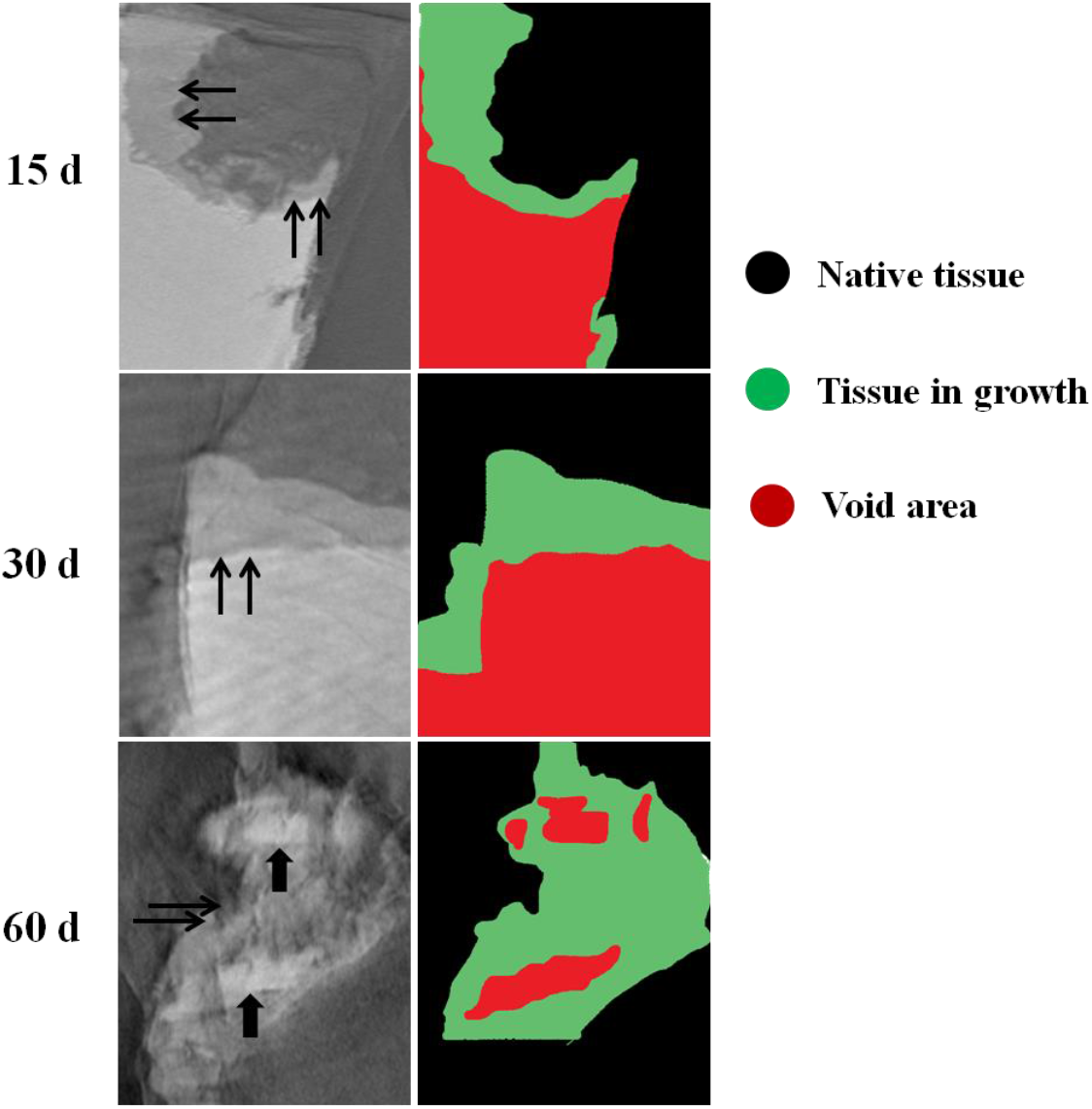
Non-invasive monitoring of tissue in growth in rabbit auricular cartilage defects evaluated by Micro-CT. Thin arrows indicates new tissue formation in defect site and thick arrows indicate void area. Almost complete healing observed 60 d post surgery.

**Fig. 13.**
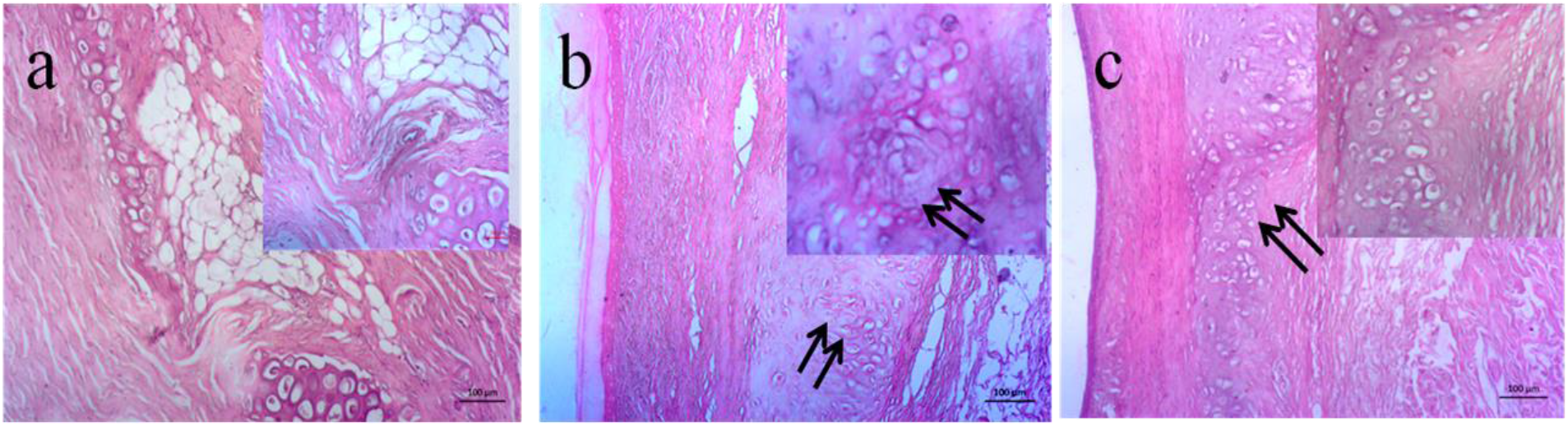
Histological analysis of rabbit auricular cartilage defect healing after 15 d, 30 d and 60 d studies. (a) 15 d studies show skin tissue growth in defect area, (b) demonstrate chondrocyte formation in defect area. Arrow indicating presence of organoid after 30 d studies and (e-f) demonstrated almost complete recovery of cartilage tissue in defect area at 60 days post-transplant studies.

#### Hematoxylin & Eosin (H&E) staining and Masson’s trichrome (MT) staining

To evaluate tissue in-growth in cartilage defect, H&E staining was performed on retrieved tissue after 15 d, 30 d and 60 d of post implantation study (Fig. 3.14). On 15th d, defect area was found to be covered with dense skin without any sign of cartilage tissue formation (Fig. 12(a)). On 30th d, chondrocytes started appearing in defect region, whereas 60 d study revealed almost complete recovery of cartilage defect (Fig. 12(b-c)). The defect region was uniformly distributed with chondrocytes, but MSs exhibited complete disappearance of the particulate morphology in course of tissue regeneration.

## 4. Discussion

Cartilage is a dense connective tissue present in all joint parts of the body enabling frictionless movement between bones. It is composed of large volume of water and mainly collagen II in the extracellular matrix. Owing to their low cellular activity and avascular nature, cartilage has limited self-repair capacity [14]. In this circumstance, tissue engineering approaches including cells, bioactive cues and engineered materials have shown immense potential towards cartilage healing. However, still some limitation has to be resolved such as biocompatibility and non-immunogenicity of the materials, less nutrient supply to deep layers of scaffold and cell sources. Considering these limitations, tissue engineered functional MS derived from *Capra* ear cartilage has immense utility to regenerate the damaged cartilage. Therefore, *Capra* ear cartilage, a bio-waste material has transformed to different size particles and assessed their potential for cell adhesion, chondrogenic differentiation and organoid formation towards cartilage defect regeneration.

The isolation of MS has several steps involving skin elimination, pulverization, decellularization and particles separation. For decellularization, NaOH treatment has performed to remove the cells. Generally, the well-established decellularization agents including SDS, Triton-X100, CHAPS etc. have several limitations like toxicity, inefficient DNA elimination. Moreover, optimal concentration of NaOH used in decellularization is important to minimize tissue distortion as well as sGAGs loss. The developed optimized process of decellularization has potential to complete removal of cells as well as DNA while preserving satisfactory concentration of sGAG in decellularized particles [15].

Collection of MS at different time points through phase separation of alcoholic dispersion has produced different sized particles (100, 150 and 300 μm MS). Particles suspended in alcohol returns back to its native state and separates in different phases according to their size, loose fibrous area and particle density. Large particles undergo settling under gravity, while small particles freely float in the liquid suspension.

The morphometric analysis of MS showed diverse characteristics of isolated particles including size, shape, circularity, loose fibrous area, dense and porous area. Such variable characteristics facilitate adequate cell adhesion on MS surface transforming them into functional micro tissues after certain period of culture. Quantitative analysis of particles morphology showed that 100 MS could preserve required properties of excellent cell adhesion and MS based organoid formation. Mean and median gray scale values of particles could be correlated with ratio between loose and dense area of particles determining foaming tendency of isolated particles during pulverization and decellularization. Nature of pore size distribution also reveals particles solidity, loose fibrous area and porous properties.

Live–dead assay results can be correlated with cell adhesion and biocompatibility properties of MS. Excessive loose fibrous network of small particles demonstrated excellent adhesion and infiltration of cells compared to other extended size particles. Therefore, cell monolayer formation was observed on small particles surface (100 MS).

The device fabricated for MS organoid culture could maintain appropriate microenvironment for extended time length of culture as compared to well-known hanging drop culture. Hanging drop culture is limited by volume of a drop cast limiting the quantity of cell-material construct to be used for organoid formation. Moreover, size of developed micro tissues was very small with loose cell-material bonding and media color changed after 3-4 d. However, the fabricated device has capacity to hold upto 500 μL media and curvature of this holder appropriately grasps the organoid for long time culture with 3D geometry.

*Ex-vivo* spheroid culture suggests that small MS has enormous potential for organoid formation within small period of time. The widely spaced fibers in small MS with thickness less than cell diameter has immense influence in improving cell adhesion, shape adaptation and superior extension towards organoid formation [17]. On the other hand, loose fibrous area with porous structure of small MS facilitated rapid cellular infiltration to the core of the particles inducing superior cell-material interaction. Therefore, hanging culture of cell-100 MS construct affords organoid formation after certain period of culture in absence of exogenous growth factor. Moreover, particles derived from native tissues contain various bioactive factors or molecular cues that stimulate ADMSCs differentiation into chondrogenic lineage. F-127 hydrogel was used to provide three-dimensional culture environment for organoid maturation.

The plausible explanation for elucidating probable mechanism of organoid formation by small particles is as well. The isolation of MS involves pulverization, decellularization and lyophilization of Capra ear cartilage tissue. Pulverization process using blender effectively tears down the small particles from its intrinsic form provides extensive fibrous network in surrounding area [18]. In native particles, fibrous network is usually formed towards the edge of particles, while central remains intact. Such structure limits diffusion of nutrients and oxygen delaying cell proliferation. However, cells are more prone to adhere on loose fibrous area either by direct contact of cell membrane to particle fiber or by intrusion of fibrous matrix directly to cell membrane near adjacent microvilli and stretches into polygonal or elongated shapes along the fiber orientation [19]. Moreover, loose fibrous area is extremely suitable for tissue remodeling and regeneration by facilitating cell infiltration into the particles. The higher surface-to-volume ratio of particles also provides more sites for cell adhesion, resulting in higher cell growth on particle surfaces (Fig. 14).

**Fig. 14.**
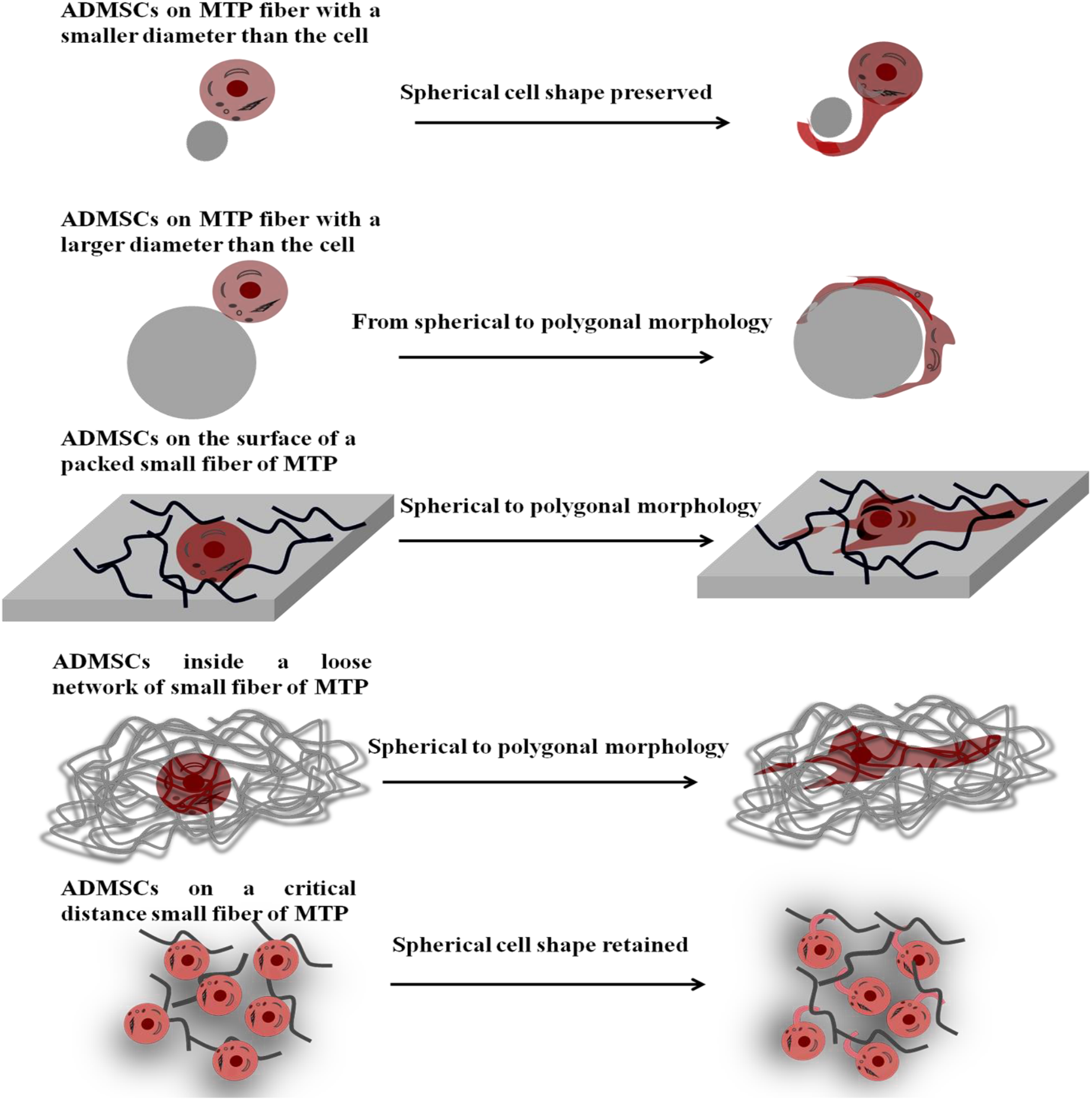
Schematic representation of plausible correlation between particle fiber size and cell morphology. Expected morphological changes of ADMSCs after seeding on MS small fiber. The small diameter of MS fiber retained the spherical morphology of cell where larger diameter preserved polygonal morphology. The small fiber of several particles close enough to each other built packed fibrous network, considered as 2D surface for transformation of spherical cell shape to polygonal. The loose fiber network with enough closeness also alters the cell morphology of spherical to polygonal. If, the fiber size enough to small than the diameter of ADMSCs and the critical distance between fibers seems to be above the cell diameter (15-20 μM), the cells retained their spherical morphology.

In hanging culture model under gravitational force, MSs gradually settle down towards bottom forming cell-particle aggregates [17]. Inter-cellular distance has a major influence on aggregate formation since larger distance between cells inhibits inter-cellular cross talk [20]. Smaller particles ensure formation of compact clusters as the distance between particles is lower than the threshold distance for cell-cell communication, and cell density on smaller particles surface is also higher. Such micro-tissue scaffolds tend to form stable aggregates [21].

Small particles with more effective surface area per gram would have more number of pendent fibers along the surface which may be responsible for favorable MS based organoid formation. The cartilage matrix related genes (COLII, SOX 9 and Aggrecan) were relatively overexpressed with small particles (100 MS) compared to large particles as evidenced by qRT-PCR results. It may be assumed that the small particles possess sufficient loose fibrous area maintaining critical distance above cell diameter and preventing polygonal chondrocyte morphology of differentiated ADMSCs. Moreover, chondrocytes could be de-differentiated into fibroblasts after certain time with a change in polygonal morphology into spindle-shapes. qRT-PCR data predicts that fibrous matrix of small particles could maintain appropriate contact with ADMSCs providing a natural chondrogenic niche for stem cells differentiation to chondrocytes. In contrary, chondrogenic related gene expression in 200 MS was relatively down regulated compared to 100 and 300 MS. To explain this finding, further investigation is required.

In summary, 100 μm MS has various superior properties including fibrous matrix, loose fibrous area, extreme cell adhesion and critical distance of fibers above the diameter of cell leading to faster organoid formation with strong cell-MS bonding. Therefore, 100 μm MS micro tissue organoid was transplanted into rabbit auricular cartilage defect for regeneration. After 60 d study, the cartilage defect was almost recovered compared to 15 and 30 d studies supported by findings of H&E and MT staining.

## 5. Conclusions

MS, especially small particles (100 μm MS) have excellent properties for tissue regeneration as well tissue remodeling. In vitro and *ex vivo*studies revealed that tissue engineered functional MS has immense potential to form organoids. Further, in-vivo transplantation of MS organoids in rabbit auricular cartilage defect resulted in almost complete recovery after 60 d. Therefore, ADMSCs infiltrated small particles have enormous potential for complete regeneration of damaged cartilage.

## Notes

### Competing Interest Statement

The authors have declared no competing interest.

### Summary of Updates

The title has been updated.

